# Long range allostery mediates the regulation of plasminogen activator inhibitor-1 by vitronectin

**DOI:** 10.1101/2022.07.19.500692

**Authors:** Kyle Kihn, Elisa Marchiori, Giovanni Spagnolli, Alberto Boldrini, Luca Terruzzi, Daniel A. Lawrence, Anne Gershenson, Pietro Faccioli, Patrick L. Wintrode

## Abstract

The serpin plasminogen activator inhibitor 1 (PAI-1) spontaneously undergoes a massive structural change from a metastable, active conformation, with a solvent accessible reactive center loop (RCL), to a stable, inactive or latent conformation in which the RCL has inserted into the central β sheet. Physiologically, conversion to the latent state is regulated by the binding of vitronectin which retards the rate of this latency transition approximately 2-fold. We investigated the effects of vitronectin on the PAI-1 latency transition using all-atom path sampling simulations in explicit solvent. In simulated latency transitions of free PAI-1, the RCL is quite mobile as is the gate, the region that impedes RCL access to the central β sheet. This mobility allows the formation of a transient salt bridge that facilitates the transition, and this finding rationalizes existing mutagenesis results. Vitronectin binding reduces RCL and gate mobility by allosterically rigidifying structural elements over 40 Å away from the binding site thus blocking the transition to the latent conformation. The effects of vitronectin are propagated by a network of dynamically correlated residues including a number of conserved sites that have previously been identified as important for PAI-1 stability. Simulations also revealed a transient pocket populated only in the vitronectin bound state which corresponds to a cryptic drug binding site identified by crystallography. Overall, these results shed new light on regulation of the PAI-1 latency transition by vitronectin and illustrate the potential of path sampling simulations for understanding functional conformational changes in proteins and for facilitating drug discovery.

## Introduction

The inhibitory serpin plasminogen activator inhibitor 1 (PAI-1), is a key protein in the negative regulation of blood clot clearance (fibrinolysis) primarily targeting tissue type plasminogen activator (tPA) and urokinase type PA (uPA) (1). Like all inhibitory serpins, PAI-1 employs the canonical “mouse trap” suicide inhibition mechanism. The trap is initiated by formation of the serine protease acyl-enzyme intermediate and proteolytic cleavage of the serpin’s reactive center loop (RCL). Upon RCL cleavage, serpins undergo a massive conformational change, in which the RCL inserts into the central β-sheet (sheet A) expanding sheet A from 5 to 6 beta strands. If RCL insertion is fast enough, the acyl-enzyme bond stays intact and the serpin pulls apart the protease active site of the protease, inactivating the enzyme, and yielding a thermodynamically stable serpin conformation (2, 3). Structural and phylogenetic studies have identified several regions in the serpin fold that are especially important in facilitating and regulating this conformational change (4–6). Two that are of particular significance for the present study are the “gate”, consisting of strands 3 and 4 of β-sheet C, and the “breach”, located at the top of strands 3 and 5 of β-sheet A (Figure 1A). The gate poses a steric barrier to movement of the C terminal portion of the RCL, and thus impedes loop insertion in the absence of proteolytic cleavage. The breach is the site of the initial separation of strands 3 and 5 in β-sheet A which allows loop insertion to begin.

**Figure 1.**
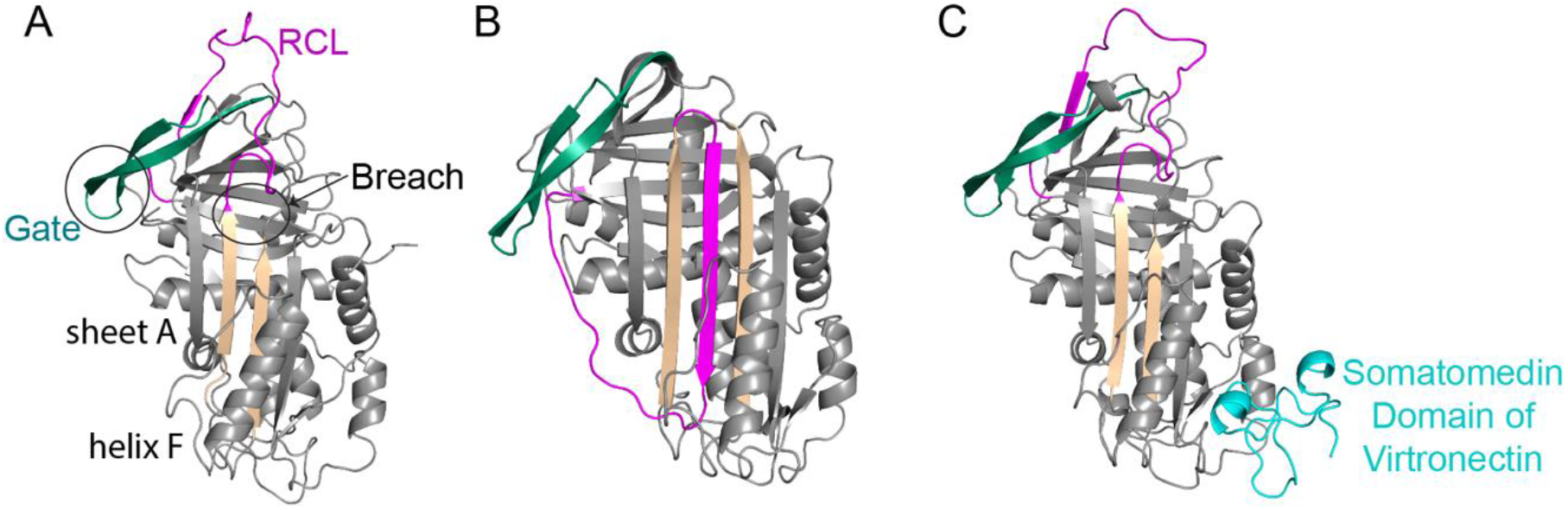
PAI-1 conformational lability. (A) The structure of active PAI-1 with a solvent accessible RCL (PDB: 3QO2 (7)). (B) The structure of latent PAI-1 (PDB: 1DVN (8)). (C) The structure of active PAI-1 in complex with the somatomedin domain of vitronectin (cyan) (PDB: 1OCO (9)). For vitronectin bound PAI-1, the missing electron density for the solvent exposed RCL was modeled using the ModLoop server (10). In all structures, the RCL is shown in magenta, the gate is shown in green, and strands 3 and 5 A are shown in tan.

PAI-1 is an unusual member of the serpin family in that RCL insertion can also occur spontaneously, without cleavage by a target protease. This spontaneous deactivation, or latency transition, occurs when the intact RCL inserts itself into sheet A, leading to a thermodynamically stable but inactive conformation of PAI-1 (Figure 1B) (5). The latency transition is one way to regulate PAI-1 activity. An additional level of regulation is achieved through binding of PAI-1 to the cell adhesion factor vitronectin (VTN) (Figure 1C) leading to an approximately 50% increase in the half-life of the active state (11). Regulation of PAI-1 activity is critical for human health, as high levels of active PAI-1 are associated with both poor prognoses for some cancers (1) and cardiovascular diseases (12, 13), and has been shown to be an indicator for prognosis in clinically severe SARS-COVID-2 infections (14). Thus, a mechanistic understanding of PAI-1 regulation has clear implications for the treatment of an array of disease states.

A number of mutations are known to alter the half-life of active PAI-1 (15–18) but because a clear mechanistic understanding of the PAI-1 latency transition and the allosteric effects of vitronectin binding is currently lacking, it has been difficult to explain the molecular mechanisms of many of these mutations. A better understanding of the latency transition could also help in identifying cryptic pockets that might be targeted for therapeutic purposes. However, while molecular dynamics (MD) simulations can aid in elucidating molecular mechanisms, the active state half-life of free PAI-1 is 1 to 2 hours under physiological conditions (19) much longer than the millisecond time scales accessible with MD. To computationally study rare events such as serpin conformational changes one can turn to a variety of enhanced sampling methods (an incomplete list of possible methods is provided in (20, 21)).

In a previous investigation of the PAI-1 latency transition (22), we employed the Dominant Reaction Pathways (DRP) enhanced path sampling technique (23). While informative, at the time the DRP method suffered some limitations including the use of implicit solvent, and the previous simulations likely overestimated concerted opening of β sheet A early in the latency transition (24) Here we apply an improved version of the earlier method, now referred to as the Bias Functional (BF) approach, which enables us to simulate the latency transition of PAI-1 in explicit solvent and to study the effects of vitronectin binding on the latency transition (25). The BF approach uses the ratchet-and-pawl MD (rMD) method (26, 27), which requires both an initial state (the PAI-1 active conformation in the presence or absence of vitronectin) and a target state (the latent conformation), to efficiently generate a trial set of latency transition trajectories. The history dependent bias (see the Experimental Procedures for details) is based on the contact map of the target state, in this case the latent conformation. In our previous studies we used all 379 amino acids in PAI-1 for the latent state target contact map, but our current investigations show that limiting this contact map to the RCL and strands 3 and 5 A reduces the constraints on the system and results in behavior more consistent with experiment. In this study we generated multiple trajectories of the PAI-1 latency transition in explicit solvent in both the presence and absence of bound vitronectin. Our results shed new light on the nature of the latency transition and elucidate the allosteric mechanism by which vitronectin extends the half-life of active PAI-1

## Results

For wild-type PAI-1, the lifetime of the active state is about 2 hours at 37°C for the free protein (19) and is increased by approximately 50% when PAI-1 is bound to vitronectin (11). Thus, the active to latent state transition is a rare event that happens on a timescale well beyond those accessible by conventional MD simulations. We therefore used the BF approach described in the Methods to simulate this transition.

### Energy landscape of the latency transition for free PAI-1

The increased computing power made possible by the adaptation of MD codes to graphical processing units (GPUs) has enabled us to generate 30 rMD trajectories of the PAI-1 active to latent transition in explicit solvent. It has been shown (28) that in the ideal limit in which the biasing force of rMD acts along the ideal reaction coordinate (namely the committor function (29)) the transition path ensemble generated by rMD samples the correct Boltzmann distribution in the transition region. Since the collective variable (CV) employed in realistic rMD simulations provides an approximation of the committor function, this scheme can only yield semi-qualitative information about the energy landscape and a lower-bound estimate of the relevant free-energy barriers.

We used our rMD trajectories to estimate the free energy landscape in the plane formed by the root mean squared displacement (RMSD) from the latent state and the fraction of contacts which are only present in the latent state, the “native” contacts Q (Figure 2). While this method does not enable us to quantitatively estimate barriers and kinetics, we expect that the presence and relative locations of metastable states along the transition pathway should be accurately predicted. Several features are evident in this landscape for free PAI-1. There are two relatively deep and one shallower minima located at high RMSD and low Q. To insert into sheet A, the RCL must circumnavigate around the so-called gate formed by strands 4C, 3C and the connecting loops. All three of the minima at low Q are populated by structures in which the RCL has not yet moved around the gate. In minimum 1, strand 1C, at the distal end of the RCL, has detached from sheet C, but there are few other structural changes from the active form. In the second minimum, the proximal end of the RCL (residues ~330-340) has moved approximately 11 Å down and across the front of β-sheet A, while the gate has been significantly displaced from its original position with the tip of the gate, defined by the turn connecting strands 4 and 3C (residues ~191-195), moved forward by 9 Å. Further displacements of both the RCL and the gate are evident in minimum 3. The structures populated in minima 2 and 3 suggest that these minima together are a good candidate for the hypothesized pre-latent state of PAI-1 (see below).

**Figure 2.**
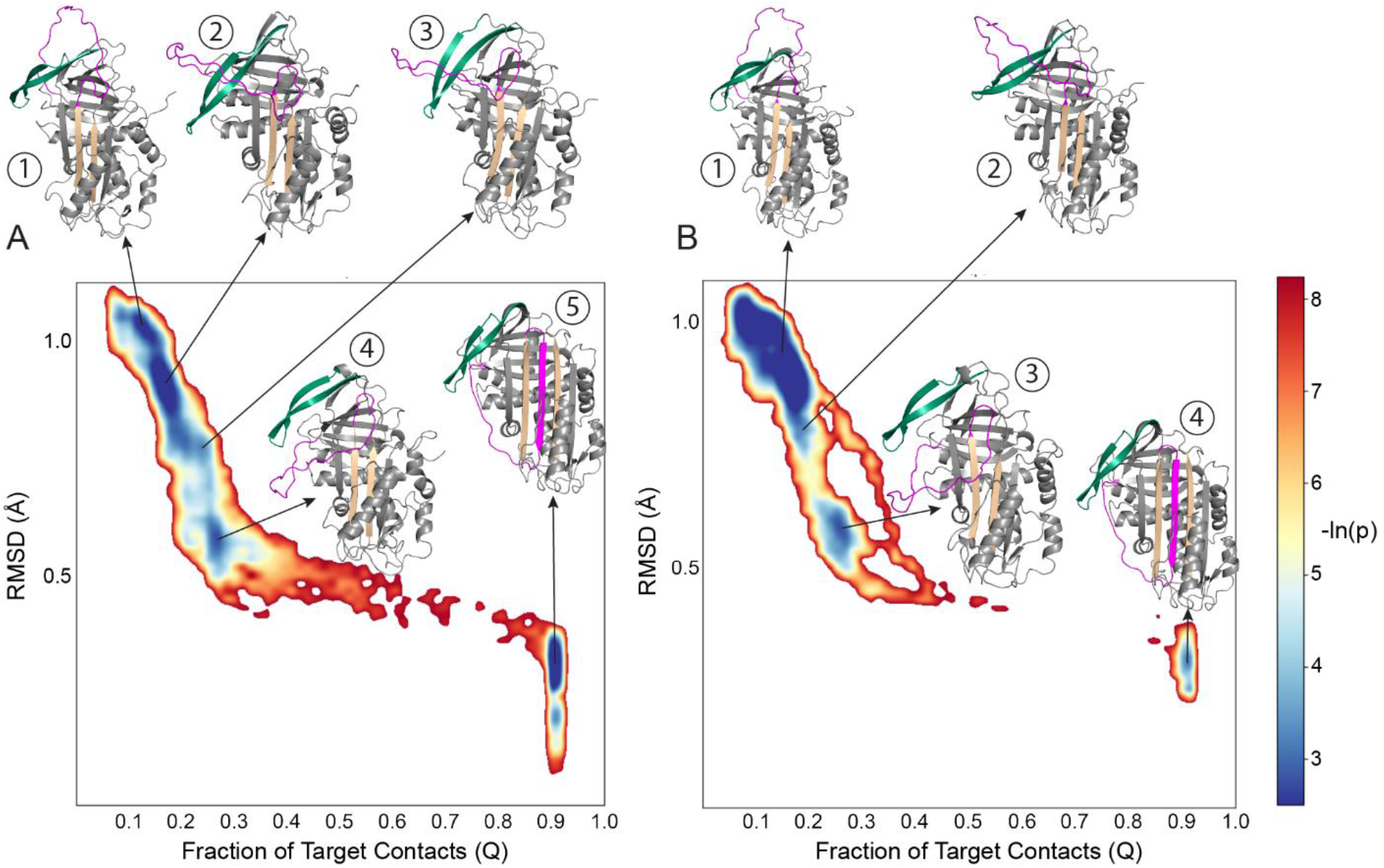
Approximate free energy landscapes for the PAI-1 latency transition. Frames are projected onto the plane defined by the RMSD from the latent structure (Y axis) and fraction of “native” contacts, contacts formed only in the latent state (X axis). (A) Simulations of PAI-1 alone (30 rMD trajectories). (B) Simulations of PAI-1 bound to the somatomedin B domain of vitronectin (20 rMD trajectories) Vitronectin is not shown for clarity. The heat map indicates approximate free energy (proportional to population density), plotted as the negative logarithm of the probability distribution of conformations (-ln(p)). Centroid structures (colored as in Figure 1) from each of the minima identified are shown. Continuous Q and RMSD were binned in a 65-by-65 bin matrix.

The fourth minimum, centered around an RMSD of 0.5 Å and a Q of 0.25, corresponds to an intermediate state in which the RCL has successfully circumnavigated the gate but has not yet fully inserted into sheet A, largely due to the barrier posed by helix F which lies in its path. This state is similar to the main long lived intermediate identified in our earlier study of the latency transition (22). However, as pointed out by Jorgensen and colleagues, in our previous simulations strands 3 and 5A were separated by ~8 Å along their entire length allowing facile insertion of the RCL, and such a large opening of sheet A is not consistent with the results of hydrogen-deuterium exchange mass spectrometry experiments probing the PAI-1 latency transitions (24). By contrast, in the current simulations where the solvent dynamics are explicitly accounted for and the contact map is limited to contacts formed between the RCL and strands 3 and 5A in the latent structure, even after the RCL passes the gate there is minimal opening of sheet A. The final stage of the transition involves the displacement of helix F and completion of RCL insertion into sheet A. As evident from Figure 2, a large free energy barrier separates minimum 4 from the final minimum that corresponds to the latent state. Nevertheless, the majority of trajectories that reach minimum 4 (23 of 30) successfully complete this transition.

In the latent state, strand 1C at the distal end of the RCL detaches from sheet C and the RCL inserts between strands 3 and 5 in the central β sheet A (8, 31). A number of studies have found experimental evidence for a so-called pre-latent state of PAI-1, in which strand 1C has detached but RCL insertion has not proceeded (30, 32, 33). PAI-1’s ability to populate this strand 1C detached state is thought to be important for the latency transition. Our previous simulations using implicit solvent identified a single long lived metastable intermediate which agreed with several experimental observations on the proposed pre-latent state, and in this proposed intermediate the RCL had already circumnavigated the gate (22). The present study has identified two additional significant long lived intermediates, minima 2 and 3, in which the RCL has not yet moved past that gate and is not even partially inserted into sheet A. Comparisons with available experimental evidence suggest that the structures populating these earlier, less perturbed intermediates, particularly those corresponding to basin 3, provide better candidates for the pre-latent state (Figure 3A). In particular, Hägglof *et al*. estimated the distances between several pairs of residues, which were mutated to cysteine and fluorescently labeled using a combination ofForster Resonance Energy Transfer (FRET). Additionally, using double cysteine mutagenesis, they showed that several pairs of residues distant in the crystal structure of active PAI-1 come close enough to form disulfide bonds. Table 1 shows that the relevant residue-residue distances measured in the centroid structure of basin 3 - but *not* in the crystal structure of active PAI-1 - are consistent with the experimental measurements of Hägglof *et al*.. The lone exception is the distance between His185 and Ser344. However, unbiased MD simulations initiated from the basin 3 structure indicate that the distance distribution of these residues is multimodal and they readily sample distances consistent with Hägglof *et al*. (Figure 3B).

**Figure 3.**
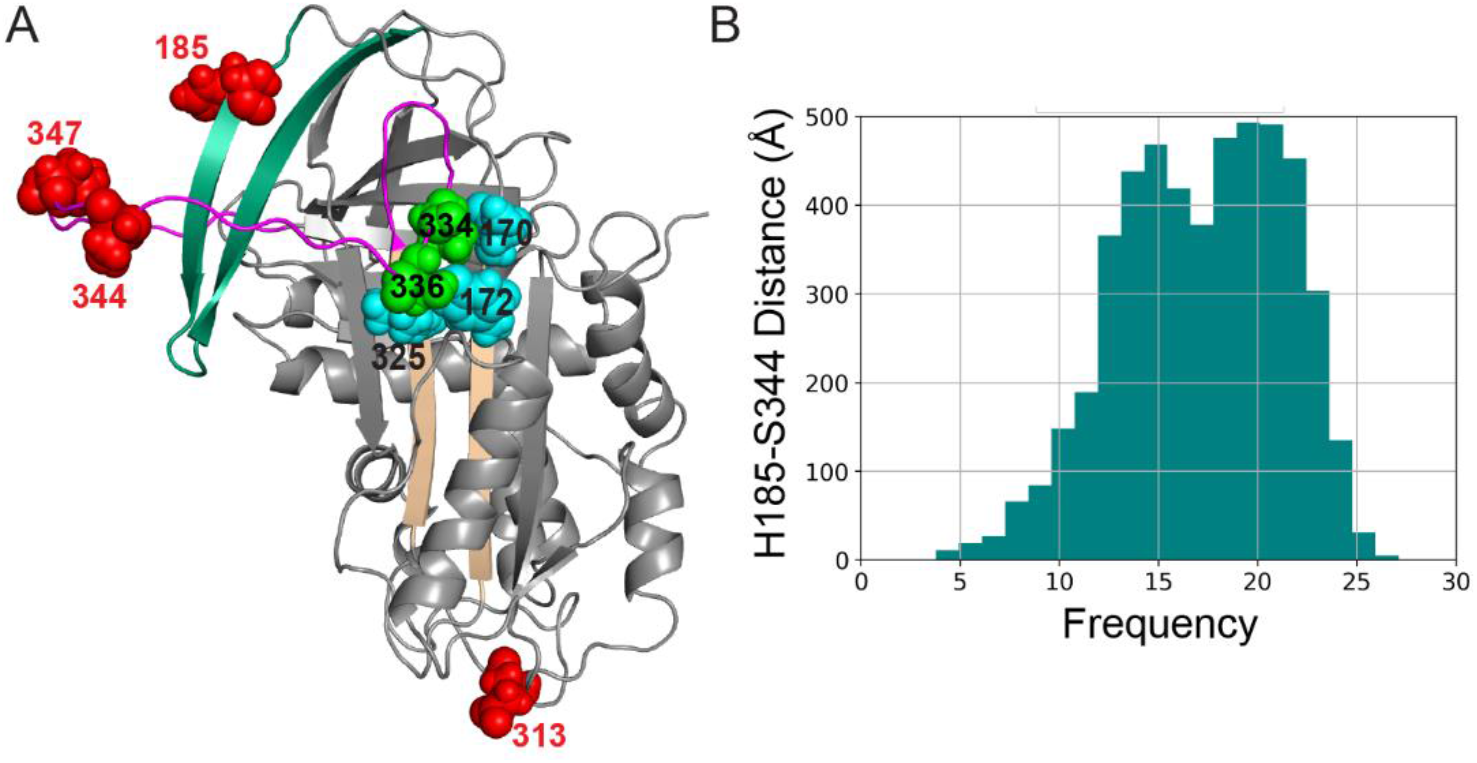
Model of pre-latent PAI-1. (A) Candidate structure for the pre-latent state of PAI-1. Residues mutated to Cys and used by Hägglof et al. to measure FRET distances(185, 313, 344, and 347) are shown as red spheres. Residues used for disulfide crosslinking experiments are shown as green spheres for residues in the RCL (334 and 336) and cyan spheres for residues in sheet A (170, 172, and 325). (B) Histogram of distances between residues 185 and 344 during unbiased MD simulations started from basin 3.

**Table 1.** Distances between selected residues in PAI-1 measured by Haägglof *et al*. (30), measured in the centroid structure from basin 3 of rMD simulations, and measured in the structure of active PAI-1.

| Distance | Haägglof *et al.* (30) | Centroid Basin 3 | Active (PDB: 3QO2) |
| --- | --- | --- | --- |
| Glu313-Met347 | $55.1 \pm 2.1$ Å | 58.4 Å | 71.5 Å |
| Glu313-Ser344 | $55.2 \pm 2.1$ Å | 55.3 Å | 73.5 Å |
| His185-Ser344 | $\leq 11.6$ Å | See Fig. 3B | 21.5 Å |
| Asn172-Ser336 | $\leq 4.5$ Å* | 4.4 Å | 8.6 Å |
| Lys325-Ser336 | $\leq 4.5$ Å* | 4.4 Å | 19.1 Å |
| Tyr170-Ser336 | $\leq 4.5$ Å* | 4.3 Å | 16.3 Å |

\*The value of $\leq 4.5$ Å is inferred from the fact that these residue pairs can form disulfide bonds when mutated to cysteines. See Materials and Methods for further details.

Although the proximity of RCL and sheet A residues is consistent with disulfide cross linking experiments (green and cyan spheres in Figure 3 A), the RCL has not inserted into sheet A. Additionally, unlike in our previous study or in suggested models (Hagglof ref I think), the RCL has not yet circumnavigated the gate.

### Transient electrostatic interactions assist in displacement of the gate

The gate, formed by β strands 4 and 3C, is the major barrier to the latency transition, and thus significant displacement of the gate from the conformations observed in active PAI-1 crystal structures is required (5, 34). Examining our successful latency transition trajectories reveals that gate displacement is likely facilitated by a transient electrostatic interaction between Arg30, located in the loop between helix A and strand 6B, and Asp193, located at the tip of the gate (Figure 4). While the oxygen-nitrogen distance averages 4.0 Å, the cutoff that is commonly used to define salt bridges in proteins (35), the distance of closest approach ranges from 5.7 to 2.4 Å in the successful transitions. Significantly, movement of the RCL around the gate corresponds to the distance of closest approach (Figure 4A). The close approach of these two residues could be a trivial consequence of being pushed by the RCL, which is acting under the influence of the rMD force. To control for the influence of the ratchet force we examined the distribution of Arg30-Asp193 distances in a 500 ns unbiased MD simulation started from the centroid structure of basin 3. In the crystal structure of active PAI-1, Arg30 and Asp193 are separated by ~10 Å. Nonetheless, the MD simulations reveal that the distribution is bimodal with a pronounced tendency to sample close distances (and this tendency is attenuated by the presence of bound vitronectin) (Figure 4B). Unbiased simulations beginning from the centroid structures of basins 1 and 2 indicate that the propensity to sample close Arg30-Asp193 distances is even more pronounced in basin 1, although not in basin 2 (Figure S1).

**Figure 4.**
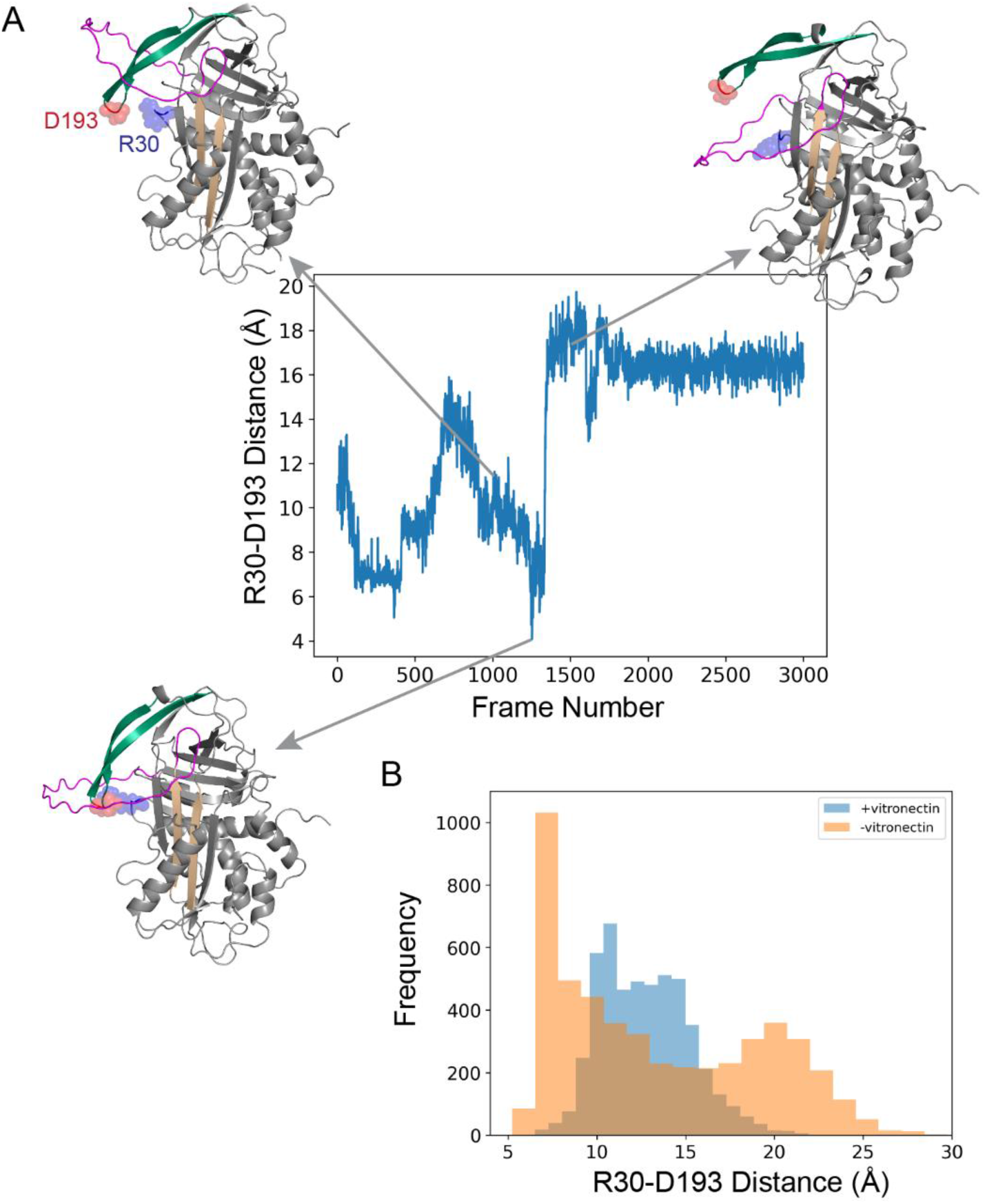
Passage of the RCL is assisted by a salt bridge (Arg30-Asp193) between the gate and the body of the serpin. (A) Distance between the Arg30(N) and Asp193(O) during an example rMD simulation of the latency transition for free PAI-1. Structures from the trajectory are shown with Arg30 and Asp 193 indicated as blue and red spheres respectively. (B) Histogram of Arg30-Asp193 distances from 500 ns unbiased simulations starting from the centroid of basin 1. The distribution for free PAI-1 is in orange and the distribution for the PAI-1:vitronectin complex is in blue.

These observations suggest that electrostatic interactions between the gate and the helix A to strand 6B turn contribute to PAI-1 function by facilitating the latency transition. We investigated this possibility further by examining the conservation of charged residues at positions 30 and 193 in the PAI-1 family. Although numerous putative PAI-1 serpins can be identified by sequence analysis, the existence and kinetics of the latency transition have been confirmed experimentally primarily in the jawed vertebrates by Jendroszek *et al*., who showed that the kinetics of the latency transition are highly conserved in PAI-1s from jawed vertebrates which share as little as 50% overall sequence identity (36). Based on this work along with studies of the evolution of the plasminogen activation system among chordates (37), we constructed sequence alignments of human PAI-1 and 16 other PAI-1s from jawed vertebrates. PAI-1, also called SerpinE1, is a member of the serpin E clade, so as a control we also aligned human PAI-1 and 16 SerpinE2 family members from the same jawed vertebrates (Figures S2 and S3).

Oppositely charged residues at positions 30 and 193 are 65% conserved in the PAI-1 alignment. By contrast, in the SerpinE2 alignment the residue corresponding to Arg30 is positively charged, either Asp or Glu, while the residue corresponding to Asp103 is a conserved aspartic acid. The change in charge at residue 30 in the SerpinE2s suggest that interactions between the gate and the body of the serpin are electrostatically disfavored In the SerpinE2s which, to our knowledge, do not readily populate the inactive latent state. For the PAI-1 alignment, if we relax our criteria to include charged residues at other positions in the helix A-strand 6B loop (which would still allow for a salt bridge to form) we find 88% conservation of a charge-charge interaction in the PAI-1 alignment. This higher degree of conservation within the PAI-1 family along with the conserved disfavorable electrostatic interaction in the SerpinE2 family suggests selective pressure at sites in the helix A-strand 6B loop and the tip of the gate, which in turn suggests that the transient salt bridge we observe in our PAI-1 rMD simulations contributes to function.

### Vitronectin binding impedes the latency transition and allosterically rigidifies the gate region

Vitronectin binding is known to significantly reduce the probability of the latency transition (11, 19). Consistent with the measured effects of vitronectin binding only 15% (3 of 20) rMD simulations of the PAI-1:vitronectin complex complete the transition in contrast to 77% (23 out of 30) of the free PAI-1 transitions. These results indicate that bound vitronectin is able to block the latency transition even in the presence of an applied rMD force favoring the transition. In the majority of PAI-1:vitronectin rMD simulations that fail to complete the latency transition, the RCL never moves past the gate region in marked contrast to the simulations for free PAI-1 where even most failed transitions are located in basin 4 where the RCL has transited the gate (Figure 2A). These results suggest that, as expected, the gate is more difficult to displace when vitronectin is bound. In contrast to free PAI-1, the approximate free-energy landscape for the PAI-1:vitronectin complex shows only a single broad minimum at high RMSD and low Q and the complex rarely leaves this initial minimum (Figure 2B). Starting from the centroid structures of this broad minimum, we performed a total of 500 ns of unbiased MD simulations and compared these simulations to that for free PAI-1 in basin 1, which is also located at high RMSD and low Q.

Comparisons of the root mean square fluctuations (RMSF) of the gate between free PAI-1 and the PAI-1:vitronectin complex indicate that the gate is less mobile in the complex (Figure 5). Rigidification of the gate in the presence of bound vitronectin can be seen in more detail by superimposing the centroid structures from basin 1 (high RMSD, low Q) and 3 (RMSD~0.5, Q~0.25). In contrast to free PAI-1 where the gate has been substantially displaced from its initial position in basin 3, for vitronectin bound PAI-1 the gate position is similar in these two basins.

**Figure 5.**
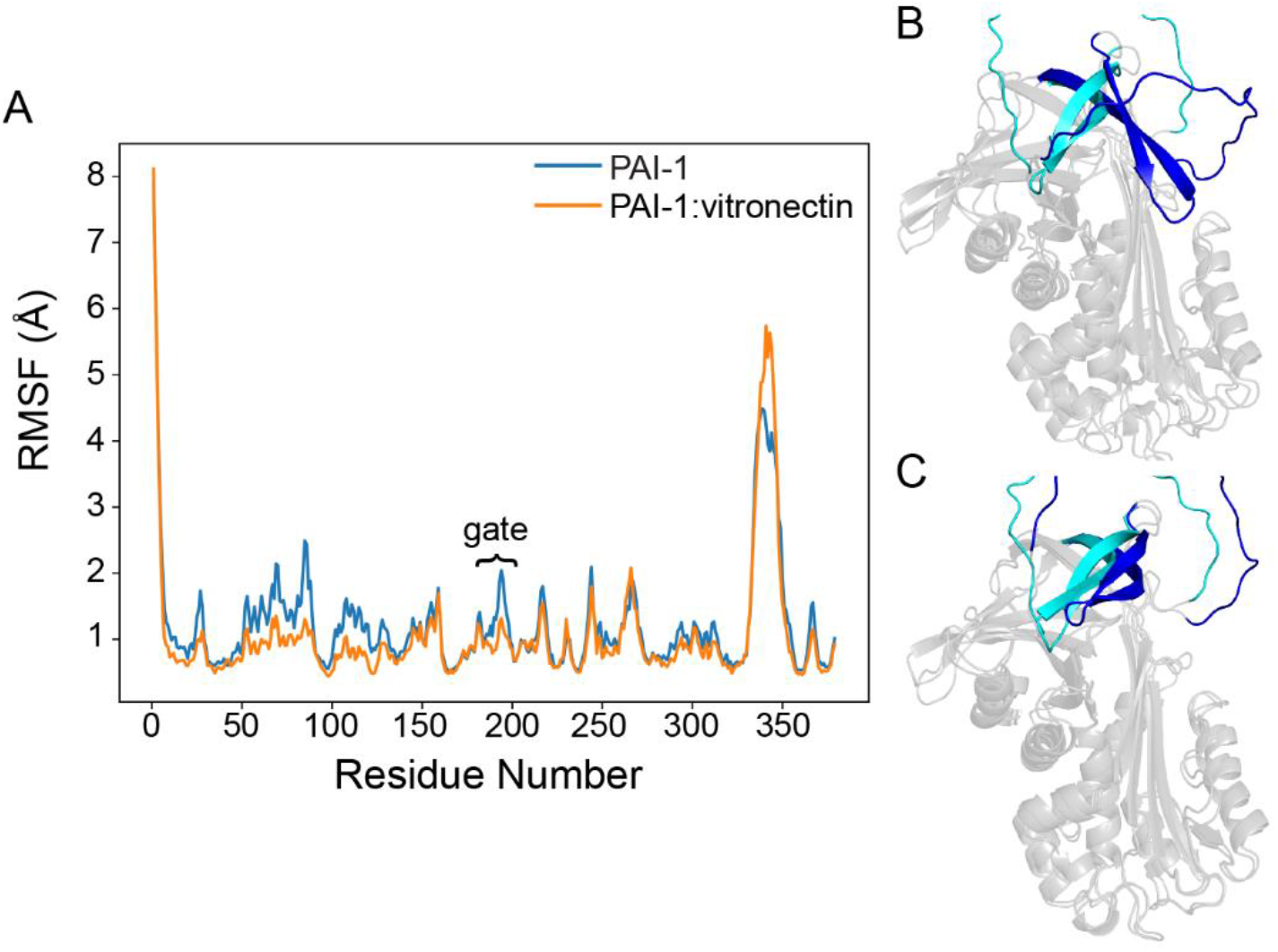
The gate is rigidified by vitronectin. (A) Root mean square fluctuations (RMSF) of alpha carbons calculated from a 500 ns unbiased MD simulations beginning from the centroid structures of basin 1 (Figure 2). Results for free PAI-1 are shown in blue and for the PAI-1:vitronectin complex in orange. (B) Superimposed structures from basins 1 (cyan) and 3 (blue) for free PAI-1. (C) Superimposed structures from basin 1 (cyan) and 2 (blue) for the PAI-1:vitronectin complex.

What is the mechanism of this rigidification? Unbiased MD simulations of PAI-1 in the presence and absence of vitronectin starting from the centroid structures of basin 1 were performed for a total of 500 ns for each state. The frequencies with which each pair of residues was in contact during the simulations was calculated, contact types were classified into salt bridges, hydrogen bonds, and van der Waals contacts using the statistical package getContacts (38), and the contact frequencies for PAI-1 in the presence and absence of vitronectin were compared.

In free PAI-1, van der Waals contacts between the RCL and strand 3C and between the tip of the gate and the body of the molecule were formed relatively frequently. These contacts formed because the RCL was bumping into the gate. In contrast, for the PAI-1:vitronectin complex, increased van der Waals contacts appear to mainly constrain the gate. Focusing our attention to the vicinity of the gate and its interface with the body of the protein, we identified a large network of van der Waals contacts that form at least 20% more frequently in the presence of vitronectin (Figure 6). The loop that connects strand 3A to the gate contains two conserved aromatic residues, Trp175 and Phe179 (5), and Trp175 was previously identified experimentally as playing an important role in regulating the stability of the active state of PAI-1 (7, 17). Interestingly, these two aromatics and Thr177 show increased van der Waals interactions with Gln204 and Asn206 in strand 4C (dark blue spheres in Figure 6). There is also an increase in interaction frequency between Phe179 and Met202, although this does not quite reach the significance cutoff of 20%. This interaction is notable because our analysis predicts that Met202 plays a major role in propagating the allosteric signal from the vitronectin binding site to the gate (see below). Additionally, there are two networks of increased van der Waals contacts in sheet C: the first of these involves residues in the gate, including the turn between β strands 4C and 3C which forms the tip of the gate, likely directly constraining the gate; the second network involves residues in strands 2 to 4C likely rigidifying the sheet itself. Increased van der Waals interactions are also observed at the top of sheet A in strands 3 and 5A possibly impeding separation of these two strands and further disfavoring RCL insertion. Overall, these results suggest that vitronectin binding induces a general increase in packing density in sheet C and the gate, which is expected to contribute to increased resistance to displacement of the gate by the RCL.

**Figure 6.**
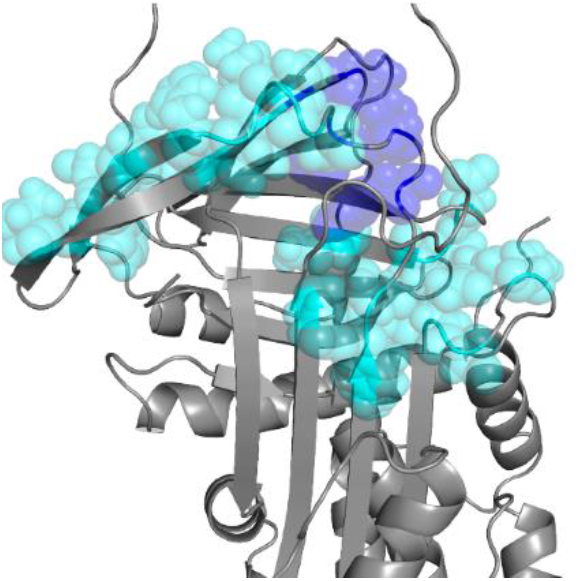
Vitronectin binding increases contacts between the gate and the body of PAI-1. Network of residues (spheres) showing increased van der Waals interactions in the presence of vitronectin. The cluster comprised of residues 175, 177, 179, 204, and 206 is shown in dark blue.

### Basis of allosteric communication between the vitronectin binding site and the gate

How does vitronectin binding influence dynamics and interactions in the gate region located 40-50 Å away? To address this question, we turned to dynamic network analysis, an established method for identifying putative pathways of allosteric communication from MD simulations which, for other systems, has successfully identified residues whose contribution to allosteric communication has been experimentally verified (39–41). This method looks for networks of residues with strongly correlated or anticorrelated motions and identifies these networks as probable pathways of allosteric communication. We examined 500 ns of unbiased MD simulations in the presence and absence of vitronectin starting from the centroid structures in basin 1 of the rMD simulations using the <u>W</u>eighted <u>I</u>mplementation of <u>S</u>uboptimal <u>P</u>aths (WISP) method (42) which requires a set of “source” and “sink” residues as inputs. To account for the effects of vitronectin binding, the source residues included all of the residues comprising the vitronectin binding site as defined by PDBsum (43). The sink residues included positions 182-205, which comprise the gate and its interface with the main body of the molecule. Since WISP by construction will always return pathways between at least one pair of source and sink residues, it is desirable to establish that the pathways identified in PAI-1 are not trivial. Two approaches were taken. In WISP the “distance” between two residues i and j is defined as −log|C_ij_|, where C_ij_ is ijth entry in the correlation matrix of residue motions calculated from an MD trajectory. The length of an allosteric pathway is then the sum of distances between pairs of adjacent residues that make up the path. Shorter path lengths indicate stronger coupling between the source and sink. For PAI-1, there is a clear shift to significantly shorter path lengths in the presence of vitronectin, indicating that these pathways are specifically coupled to vitronectin binding (Figure 7A). As a second test, WISP was run using the same source residues in the vitronectin binding site and a set of 23 sink residues located in helices G and H (Figure 7B). Both in terms of physical distance and number of intervening residues, these sink residues are roughly the same distance from the source residues as those in the gate. Not only are path lengths significantly longer on average for these sink residues, but there is no significant change in the path length distribution upon vitronectin binding.

**Figure 7.**
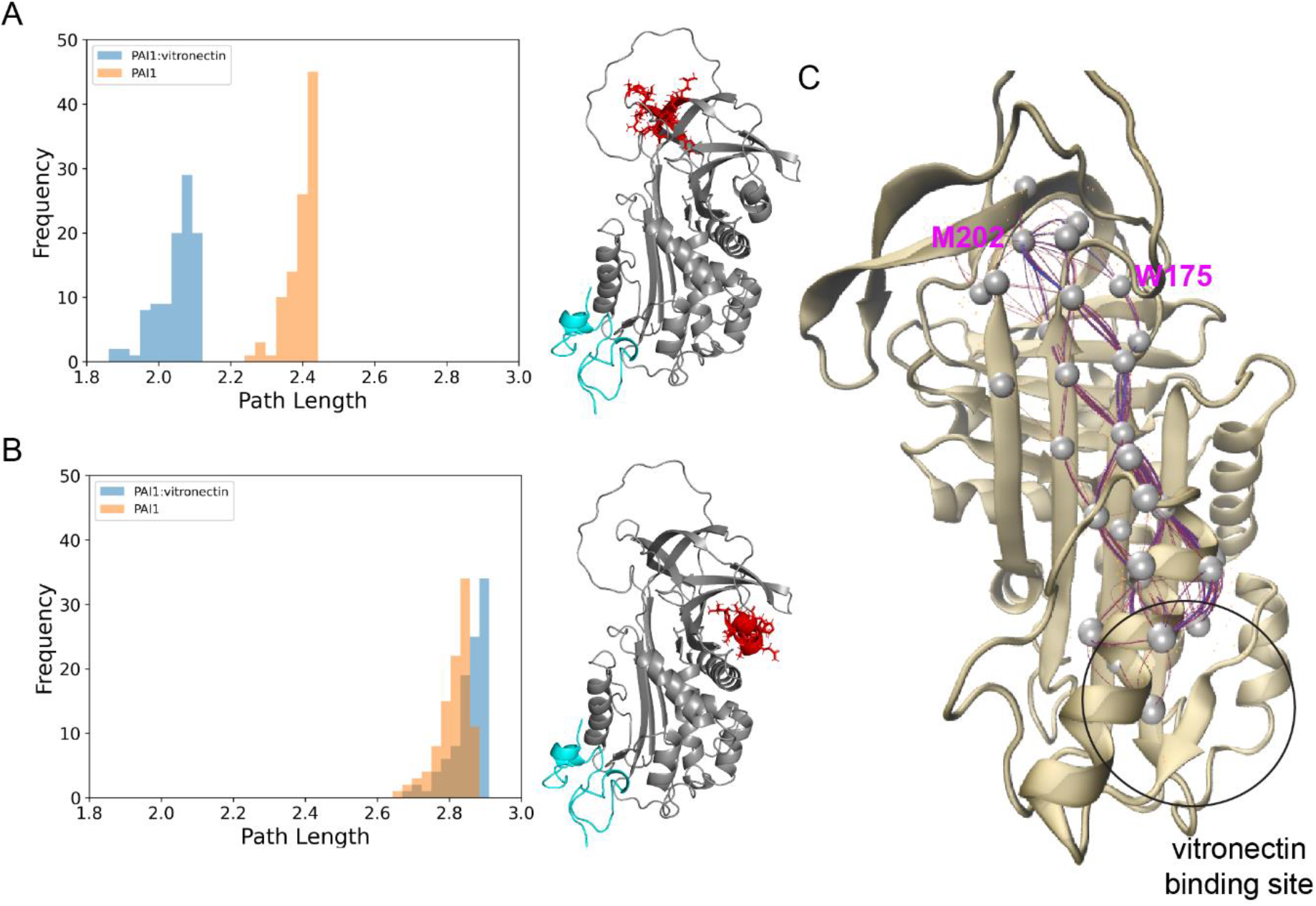
Allosteric pathways connect the vitronectin binding site to the gate. (A) Distribution of path lengths in the presence (blue) and absence (orange) of vitronectin. (B) Same as A, except sink residues were chosen in helices G and H. Locations of vitronectin (cyan) and sink residues (red) are shown on the structure of PAI-1. (C) Allosteric communication pathways between the vitronectin binding site and the gate identified by WISP (42). The blue line shows the shortest path, magenta lines show additional suboptimal paths.

The 100 most strongly connected pathways were calculated for the allosteric network between the vitronectin binding site and the gate (Figure 7C). It is clear that a relatively small number of residues have very high pathway degeneracy, playing roles in the majority of the computed pathways. These include Thr144, Tyr170, Asn172, Asn329 and Met202. Met202 is particularly noteworthy as it acts as the major “receiver” residue in the gate and is evolutionarily conserved in serpins (5).

### An allosteric pocket is formed in the presence of vitronectin

Because PAI-1 activity is associated with a number of pathogenic conditions, controlling the latency transition with small molecule inhibitors is a very active area of research (1, 12). Do the latency transition simulations reveal cryptic pockets that might be targeted for drug discovery? Crystal structures of PAI-1 and PAI-1:vitronectin, and centroid structures from each of the basins for simulations performed in both the presence and absence of vitronectin were submitted to the PASSer web server, which applies a machine learning based method to identify potential allosteric sites in protein structures (44). One site in particular was the top scoring site in structures taken from basins 1 and 2 of the PAI-1:vitronectin simulations (both basins occur before the RCL has moved around the gate), but was not detected in crystal structures of PAI-1 or PAI-1:vitronectin nor in any of the other basins from the simulations. This cryptic site is a pocket at the interface of β sheets B and C. Several of the residues that comprise this pocket, including Trp175, Thr177, Phe179, Gln204 and Asn206 were also identified by us as having van der Waals interactions that were significantly altered by vitronectin binding (see above). Additionally, Trp175, Thr177 and Phe179 were identified by WISP as part of the allosteric network connecting the vitronectin binding site to the gate.

A pocket very similar to identified by PASSer was previously identified in a co-crystal structure of PAI-1 bound to a small molecule inhibitor that binds reversibly and allosterically inhibits interactions both with proteases and with vitronectin (45) (Figure 8). Interestingly, while drug binding to this site *prevents* binding of vitronectin, in our simulations this pocket is observed only in the *presence* of bound vitronectin. Comparison of the drug bound crystal structure and the PASSer pocket reveals that while most of the same residues are involved in both the experimental drug binding pocket and our proposed allosteric pocket, the side chain arrangements are not identical. It is possible that subtle alterations of structure in this region upon drug binding are propagated to the vitronectin binding site through the same allosteric network identified from our simulations leading to vitronectin dissociation.

**Figure 8.**
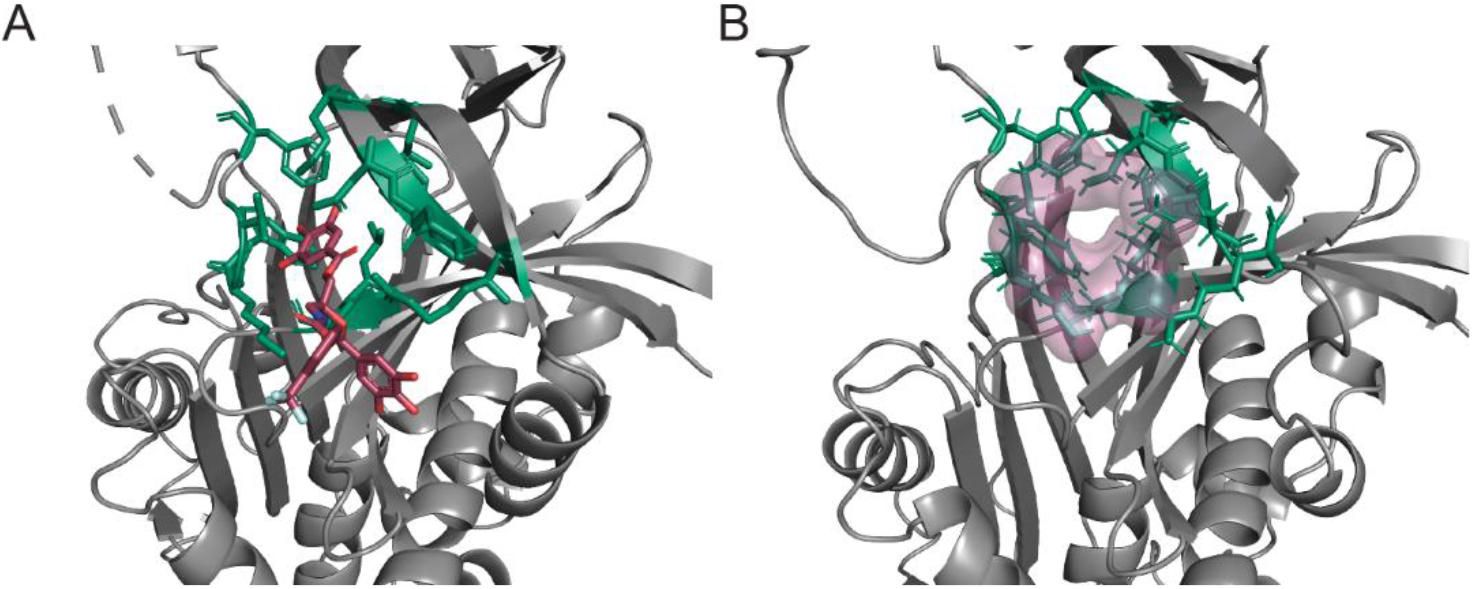
Allosteric pocket formed in the presence of vitronectin. (A) Crystal structure of PAI-1 complexed with a small molecule inhibitor (PDB: 4G80 (45)). The small molecule is shown as blue sticks. (B) Allosteric pocket detected in structures from basins 1 and 2 in the PAI-1:vitronectin simulations. The pocket surface is shown in red. In both A and B, residues 175, 176, 179, 204, 206, 207, 208, 226, 268, and 273 are shown as green sticks.

## Discussion

Serpins face a unique challenge in balancing the stability required to maintain their active conformation with the plasticity required for the massive conformational change they must undergo as part of their function. This balance is especially critical for PAI-1 because spontaneous inactivation via conversion to the latent state is a part of its functional regulation. The present study sheds additional light on this balance, bringing particular attention to the key role played by the flexibility of the gate region in regulating PAI-1 stability. The conformational lability of the active form of PAI-1 was highlighted by the discovery of evidence for a pre-latent state. Characterization of this pre-latent form of PAI-1 has, however, focused almost entirely on the conformation of the RCL (30, 32). Our simulations of the latency transition have identified a metastable conformation of PAI-1 in which the position of the RCL is consistent with all available experimental data on the pre-latent state while also highlighting the movement of the gate in adopting this structure. Unbiased MD simulations beginning from basin 1 in the free PAI-1 approximate energy landscape (Figure 2A) reveal that the gate is highly mobile even in the absence of ratchet forces driving the latency transition. The gate in these simulations frequently samples conformations consistent with passage of the RCL (Figure 4B). Simulations suggest that gate conformations that would enable passage of the RCL are stabilized by a transiently formed salt bridge between Arg30 and Asp193 that is not present in crystal structures of either the active or latent forms of PAI-1, and sequence analysis indicates that this salt bridge is conserved at least in the jawed vertebrates in which the existence of the latency transition has been experimentally confirmed. Interestingly, Haynes et al. found that eliminating this salt bridge by substituting Arg at position 30 with an uncharged amino acid extends the lifetime of the active form of PAI-1 providing experimental evidence for the proposed role of this salt bridge (18) while Lawrence and co-workers showed that mutating this arginine to glutamic acid prolonged the active half-life of PAI-1 (16). In addition, in SerpinE2s from jawed vertebrates, which to our knowledge do not readily go latent, the residue corresponding to Arg30 is positively charged (either aspartic or glutamic acid) while Asp193 is conserved (Figure S3) providing a possible electrostatic barrier to the interactions between the gate and the body of the serpin observed in PAI-1, In addition to shedding light on the latency transition, this result highlights the ability of enhanced path sampling simulations to identify transient functional interactions that may not be inferable from crystal structures.

It is worth noting that one aspect in which our simulations do not agree with experiment is the observation of slow cooperative unfolding in portions of helices A, B, and C accompanying the latency transition. Such unfolding was observed by Trelle et al. (46) using hydrogen deuterium exchange and occurred on the same timescale as the latency transition. Given the very slow timescale of these unfolding reactions (hours), it is not surprising that they were not observed in our simulations, particularly since these helices are not included in those regions of the contact map to which biasing forces were applied. However, we note that in addition to the gate, the C terminal portion of helix A and most of helix C showed significantly reduced mobility in the presence of bound vitronectin (Figure 5A). Thus, simulations suggest that mobility in these regions may be important for the latency transition and that this mobility is suppressed by vitronectin.

A major advance in understanding the regulation of PAI-1 by vitronectin was the determination of the structure of active PAI-1 in complex with the somatomedin domain of vitronectin (9). The structure revealed that the somatomedin domain binds to a region encompassing parts of helix E, strand 1A and helix F. Mutagenesis experiments subsequently identified a binding site for the non-somatomedin portion of vitronectin on helices D and E, although binding to this site was not associated with retardation of the latency transition (47). The PAI-1:somatomedin domain structure immediately suggested a mechanism by which vitronectin binding could block the transition to the latent state: the latency transition requires the movement of strands 1 and 2 A into the gap between helices E and F, and the bound somatomedin domain sterically blocks this movement. However, this mechanism, while important, still leaves some aspects of vitronectin’s effects on PAI-1 unexplained. Vitronectin binding does not just slow the transition to the latent state, it extends the lifetime of *active* PAI-1 (11). As seen for example in antithrombin and antichymotrypsin, serpins can adopt a variety of inactive, partially RCL inserted conformations that do not require displacement of the bottom of strands 1 and 2 A (48, 49). Additionally, vitronectin binding has been shown to significantly increase the inhibitory activity of PAI-1 towards both activated protein C and thrombin (50). These observations imply that vitronectin binding somehow influences structure and/or dynamics at the top of PAI-1 in the vicinity of the RCL.

The present study identifies allosteric rigidification of the gate region as a likely mechanism for the long-range action of vitronectin binding. This rigidification of the gate region rationalizes several observations on the effects of vitronectin binding on PAI-1. Due to gate rigidification, the RCL populates more solvent accessible conformations than those seen in the putative pre-latent state populated in free PAI-1 in which the “breach” region - generally thought to be the initial site for insertion - is largely blocked by the proximal residues of the RCL. RCL solvent accessibility is likely to favor protease binding consistent with the observation, noted above, that vitronectin-bound PAI-1 is more active. It has also been found that the incorporation of RCL-mimicking peptides into sheet A is accelerated by vitronectin (51). Disfavoring of pre-latent conformations due to vitronectin binding would allow RCL-mimicking peptides easier access to the breach at the top of sheet A.

The effects of vitronectin binding appear to be allosterically propagated from the binding site to the breach via a network of dynamically coupled residues. This allosteric perturbation increases van der Waals interactions in both the breach and the gate, In the breach this manifests as strengthened interactions between strand 3B and the tops of strands 3A and 5A. While strengthening interactions in sheet C both between strands 3C and 4C and with the neighboring strand 2C serve to rigidify the gate. Additional increased interactions between sheet C and strands 3A and 3B serve to strengthen the connection between the gate and rest of the molecule.

Interestingly, the conserved residue Trp175 appears in both the allosteric network identified by dynamic network analysis and in the set of residues showing increased van der Waals interactions in the presence of vitronectin. Also, Trp175 was previously identified by mutagenesis as playing a key role in modulating the stability of active PAI-1 (7, 17). Another residue of interest is Leu169 which appears in the network of residues we identify as responsible for propagating the effects of vitronectin binding. In a deep scanning mutagenesis study aimed at identifying sites that modulate PAI-1 stability, Haynes et al. found that Leu169 was the residue most enriched in missense mutations upon screening for increased stability of active PAI-1 (18). In fact, the regions identified in the deep scanning mutagenesis study of Haynes et al. as most enriched in stabilizing mutations strikingly overlap with the regions we identify as containing residues that are allosterically effected by vitronectin binding: commonalities include the top of helix F, the loop connecting helix D to strand 2A, the top halves of strands 3A and 5A, strand 3B, and strands 3C and C in the gate (18). Local regions in a protein structure may be perturbed either by mutations or by structural or dynamical changes induced allosterically. The fact that deep scanning mutagenesis and pathway sampling MD simulations have converged on common regions in PAI-1 suggests that these two studies are identifying a common mechanism for modulating the lifetime of active PAI-1. Further investigation of this mechanism throughsimulations of the newly identified stabilized PAI-1 mutants is a promising future direction.

An additional promising area for further investigation is the detection of potentially druggable pockets that are transiently formed during the latency transition. The pocket presented in this work is not evident in either active or latent PAI-1 crystal structures in the absence of bound drugbut was readily identified in structures populating basin 1 of rMD simulations carried out in the presence of vitronectin. This pocket is of particular interest as it has been experimentally verified, but additional pockets are present in other basins and in the rMD simulations of free PAI-1. It has already been shown in the context of protein folding that rMD simulations can help identify transient druggable sites not seen in crystal structures (52, 53). The present work suggests that this approach may be extended to pockets transiently formed during large scale conformational changes.

## Methods

### PAI1 Structure Generation

The crystal structure of active PAI-1 (3QO2) (7) was used with mutations introduced so the sequence matches that of wild type human latent PAI-1. The target latent structure in the rMD simulations was wild type latent PAI-1 (1DVN) (8).

### PAI-1 Vitronectin Structure Generation

The structure of PAI-1 bound to the vitronectin somatomedin B domain(PAI1:vitronectin) was generated using the x-ray crystal structure of plasminogen activator inhibitor-1 in complex with somatomedin B domain of vitronectin (PDB:10C0) (9). PAI-1 chain mutations (His150Asp, Thr154Lys, Ile354Met, and Leu319Gln) were made using the CHARMM-GUI (54) in order to match the mature wild-type (UniProt P05121) human sequence. Missing residues in PAI-1 (Val1, His2, His3, Pro4, Pro5, Ser338, Thr339, Ala340, Val341, Ile342, Val343, Ser344, Ala 345, Arg346, and Met347) were also modeled using the CHARMM-GUI.

Missing somatomedin B domain residues (Asp1 and Gln2) were modeled in using the CHARMM GUI. Sulfur-Sulfur distances in the somatomedin B domain were checked using PyMol (55) in order to ensure disulfide bonds remained intact (Cys5-Cys21, Cys9-Cys39, Cys19-Cys32, and Cys25-Cys31). PDBSum (43) was then used to ensure the binding interface of vitronectin to PAI-1 was appropriate. The resulting structure was then used in all MD simulations.

### Biased Simulations

As discussed in the introduction, in the BF approach variational filtering is applied to a set of trial transition pathways generated by rMD.

In an rMD simulation (26, 27), the rate of successful transitions across free large energy barriers is accelerated by introducing a time-dependent biasing force that depends on a collective variable, CV, z that switches on only when the system attempts to backtrack toward the reactant state, i.e.

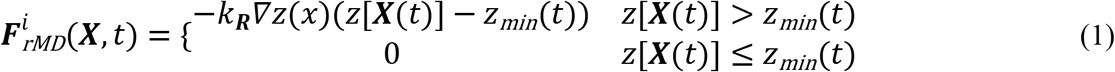

Here, **X(t)** denotes the molecular configuration at time *t*, z(X) is the biasing CV that represents a proxy of the reaction coordinate, and z*_min_*(t) is the minimum value of z(X) attained up to time *t*. A choice of CV that is commonly adopted in rMD simulations of large protein conformational transitions and folding is one that measures the overlap between the instantaneous and target state contact map:

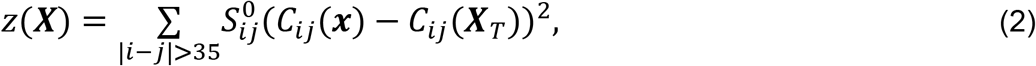

In this equation, ***X_T_*** is the target configuration (in this case, the PAI-1 latent state) and *C_ij_*(***X***) is defined as

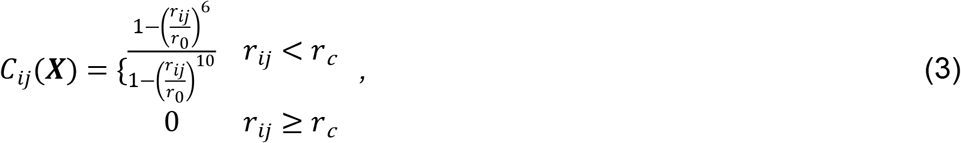

where r_ij_ is the distance between the *i*-th and *j*-th atom, *r*_0_= 0.75 nm is a reference distance for contact formation, and *r_c_*=1.23 nm is a cut-off distance.

*S^0^_ij_* in Eq. (2) is a sparse binary matrix, which can be introduced to restrict the summation to a suitable subset of atoms. In our previous work (22) we had employed a definition of CV based on the contact map for all of the atoms in PAI-1, i.e. setting *S^0^_ij_=1* for all *i,j*. We note that this choice introduces a small biasing force on atoms in regions that do not undergo structural rearrangements. While these biasing forces are generally weak, in the serpin latency transition the number of atoms that do not undergo a major structural rearrangement is quite large. As a result, the cumulative effect of these weak biasing forces might introduce a significant systematic error. We therefore decided to adopt a different choice of *S^0^_ij_* to effectively restrict the summation in Eq. (2) to the atoms of the residues in the RCL and of the residues in β-strands 3 and 5 A.

<u>T</u>he strength of the rMD basing force is controlled by the coupling constant *k**R***. Beginning with a trial value for this parameter, rMD simulations of the latency transition for free PAI-1 were repeated at successively lower values until the minimum value that consistently resulted in successful transitions to the latent state was identified. This value of *k***_R_** = 1.25×10^−4^ kJ/mol was employed for all subsequent rMD simulations of both free PAI-1 and PAI-1 bound to the somatomedin B domain of vitronectin (PAI-1:vitronectin)

In the BF approach, the set of trial trajectories generated by rMD is further processed to identify those that have the largest probability to occur in an unbiased Langevin equation. In particular, these so-called Least-Biased Trajectories (LBTs) are those with a minimum value of the functional

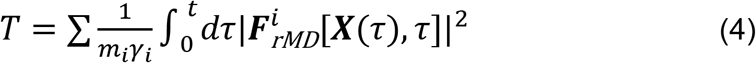

In this equation, *γ_i_* and *m_i_* are, respectively, the viscosity coefficient and mass of the *i*-th atom.

The BF scheme has been extensively validated using plain MD protein folding simulations performed on the Anton supercomputer (23, 25), against the results of deuterium-exchange experiments (56),time-resolved single-molecule FRET (57), and near-UV CD measurements (58). The BF scheme has also beensuccessfully applied to investigate amylodogenic processes (59) and in drug discovery, to identify small molecules that can interfere with the protein folding process (52). For each active conformation, a set of 20 to 30 latency transition pathways were generated by rMD and then scored according to the functional in Eq. (4).

The full ensemble of rMD trajectories was used to estimate the free energy landscape as a function of suitable pair of CVs (see the Results section), while the representative set of molecular configurations in the metastable states (to be further analyzed for cryptic pocket detection) were selected from the LBTs.

Simulations for each of the initial conditions was performed using Gromacs 4.6.5 (60) and patched with the plugin for collective variable analysis Plumed 2.0.2 (61). Each initial condition was positioned in a dodecahedral box with 15 Å minimum distance from the walls solvated with TIP3P water model (62) and 0.15 M KCl, energy minimized using the steepest descent algorithm and then equilibrated first in the NVT ensemble (using the Nosé-Hoover thermostat at 350 K, τT = 1 ps) and then in the NPT ensemble (using the Nosé-Hoover thermostat at 350 K, τT = 1 ps, and the Parrinello-Rahman barostat at 1 bar, τP = 2 ps). 20 to 30 trajectories were generated for each state by employing the rMD algorithm in the NPT ensemble (350 K, 1 bar). Each trajectory consists in 1.5 × 10^6^ rMD steps generated with a leap-frog integrator with time-step of 2 fs. Frames were saved every 5 × 10^2^ steps. The ratchet constant kr was set to 1.25 × 10^−4^ kJ/mol. Non-bonded interactions were treated as follow: Van-der-Waals and Coulomb cutoff was set to 16 Å, whereas Particle Mesh Ewald was employed for long-range electrostatics.

### Intermediate Identification

Identification of fintermediates in the latency transition was done by plotting the free energy landscape of RMSD vs Q for all trajectories in each state. RMSD was computed using Gromacs (60) while the fraction of native contacts (Q) was computed using VMD 1.9.2 (63). The energy landscape for each state was determined using a lower-bound approximation of G(Q, RMSD). This was generated by plotting the negative logarithm of the 2D probability distribution of the collective variables Q and RMSD obtained from all of the rMD trajectories. Conformations belonging to the intermediate state were clustered using the k-means approach. Distances between conformations were defined by computing the α-carbon contact maps *C_ij_(x)* (see Eq. 3) with *r_0_* = 0.75 nm. And *r_cut_* = 1.23 nm. Also, the minimum distance between *I* and *j* residues indices was set to ≥ 3 positions, to exclude residues adjacent in the sequence. One representative structure for each cluster was selected by computing the average contact map of the conformations within that cluster and then extracting the structure minimizing the distance D between its contact map and the average contact map:

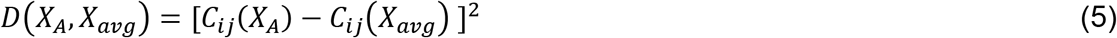

Representative structures from each intermediate were then used for unbiased conventional MD simulations.

### Intermediate conventional MD Simulations

Five repeat MD simulations for free PAI-1 and PAI-1:vitronectin intermediate structures were performed using OpenMM v7.5 (64) as follows. Inputs for the MD simulations were generated using the CHARMM-GUI (54) Input Generator web application. Simulations were carried out using OpenMM with the CHARMM36m additive force field and the TIP3P water model (65). The system was solvated in a periodic water box containing 0.15 M KCl, with box boundaries no closer than 1 nm to any solute atom. For the Lennard-Jones interaction calculations, a force switching function was applied over the range from 1.0 nm to 1.2 nm. The particle-mesh Ewald approach, with an Ewald error tolerance of 0.0005, was used for the calculation of long-range electrostatic interactions. A 2 fs time step was used for integration with temperature and pressure held constant at 298.15 K and 1 atm, respectively. Temperature was maintained at 298.15 K using a Langevin thermostat with a friction coefficient of 1 ps^−1^. Pressure was isotropically held constant at 1 bar using a Monte Carlo barostat with a pressure coupling frequency of 2 ps. Before the production run, the system energy was minimized using the L-BGFS method where 5000 steps of minimization were performed and a convergence tolerance of 100 kJ/mol was utilized. The system was then equilibrated for 125 ps in the NVT ensemble using a 1 fs timestep. During both minimization and equilibration, positional restraints were applied to the protein’s backbone and side chain atoms with a force constant of 400 and 40 kJ/mol/Å^2^, respectively. For production runs, each model and the crystallographic form were simulated for 100 ns, with structural coordinates written to the trajectory every picosecond of simulated time, resulting in 50,000 frames total. For each intermediate the five trajectories were combined and every 50^th^ frame selected to yield final trajectories of 5,000 frames. An average RMSF per residue over the course of each resulting intermediate trajectory was calculated using the MDAnalysis python package (66). Distances and distance distributions were calculated using MDTraj (67). For comparison with the data of Haägglof *et al*. on the pre-latent state, distances between the β-carbons of residue pairs 170-336, 172-336 and 172-325 were measured. Distances between the γ-carbon of residue 344 and the β-carbon of residues 313 and 185 were calculated, and the distance between the β-carbon of residue 313 and the SO of residue 347 was calculated. Distances were measured using PyMol (55). For residue pairs capable of forming disulfide bonds when mutated to cysteines, a distance between β-carbons of ≤ 4.5 Angstroms was inferred based on the analysis of Hazes and Dijkstra (68).

### Allosteric Pathway Calculations

Allosteric pathways between the vitronectin binding site and the gate were calculated from a1 μs unbiased MD simulations started from the centroid structure of basin 1 for PAI-1 +/- vitronectin. The WISP method (42) was used. Arg101, Met110, Pro111, Phe114, Thr120, Lys122, Gln123, Val124, Asp125, Arg131, Ile135, Asp138, and Trp139, were used as source residues and Leu247, Ser248, Ala249, Leu250, Thr251, Asn252, Ile253, Leu254, Ser255, Ala256, Gln257, Leu258, Ile259, Ser260, and His261 were used as sink residues. Control sink residues outside the gate were Leu247, Ser248, Ala249, Leu250, Thr251, Asn252, Ile253, Leu254, Ser255, Ala256, Gln257, Leu258, Ile259, Ser260, and His261. Resulting pathways were visualized using VMD (63).

### Contact Analysis

Contact analysis was performed using the getContacts python package (38) using the intermediate conventional MD trajectories.

### Sequence Analysis

Sequence selection was guided by work by Chana-Muñoz, et al. on the evolution of the plasminogen activation system (37). Based on their work we included at least one PAI-1 sequence from mammals, birds, crocodilians, turtles, lizards, snakes, amphibians, lungfish, coelacanths, ray-finned fishes and cartilaginous fishes. We made sure to include all of the sequences for which the active PAI-1 lifetime has been determined experimentally: human (*Homo sapiens*) (19), mouse (*Mus musculus*) (69), African lungfish (*Protopterus annectens*) (36), spiny dogfish (*Squalus acanthias*) (36) and zebrafish (*Danio rerio*) (70). We cross-referenced sequences from Chana-Muñoz, *et al*. using BLAST to locate the sequences in UniProt (71) and NCBI, with the exception of the PAI-1 sequence for African lungfish, for cases where we could not find the exact sequence, we substituted the closest database sequence as determined by E-value. Signal sequences were identified either from the UniProt annotation or using SignalP 6.0 (72).. The selected, mature (no signal sequence) sequences were aligned using Clustal Omega (73), visualized using Jalview (74) and annotated with the human PAI-1 structure (3QO2 (7)) using ESpript 3.0 (75).

### Intermediate Analysis Pocket Identification

Centroid structures from all basins for both PAI-1 and PAI-1:vitronectin simulations were submitted to the PASSer allosteric pocket detection web server using the default input parameters (44). crystal X-ray crystal structures of active, free PAI-1 (PDB: 3QO2 (7)) and PAI-1 in complex with vitronectin (PDB: 1OCO (9)) were also submitted to PASSer.

## Disclaimer & Conflict of Interest Statement

This article was prepared while Anne Gershenson was associated with the University of Massachusetts Amherst. The opinions expressed in this article are the author’s own and do not reflect the view of the National Institutes of Health, the Department of Health and Human Services, or the United States government. PF and GS are co-founders of Sibylla Biotech (www.sibyllabiotech.it), a company involved in early stage drug discovery.

## Acknowledgements

We thank Laura M. Haynes, Zachary M. Huttinger, Andrew Yee, Colin A. Kretz, David R. Siemieniak, Daniel A. Lawrence, and David Ginsburg for access to their PAI-1 deep mutational scanning data and results prior to publication. We also thank Emiliano Biasini for helpful discussions.

## Supplementary Materials

**Figure S1.**
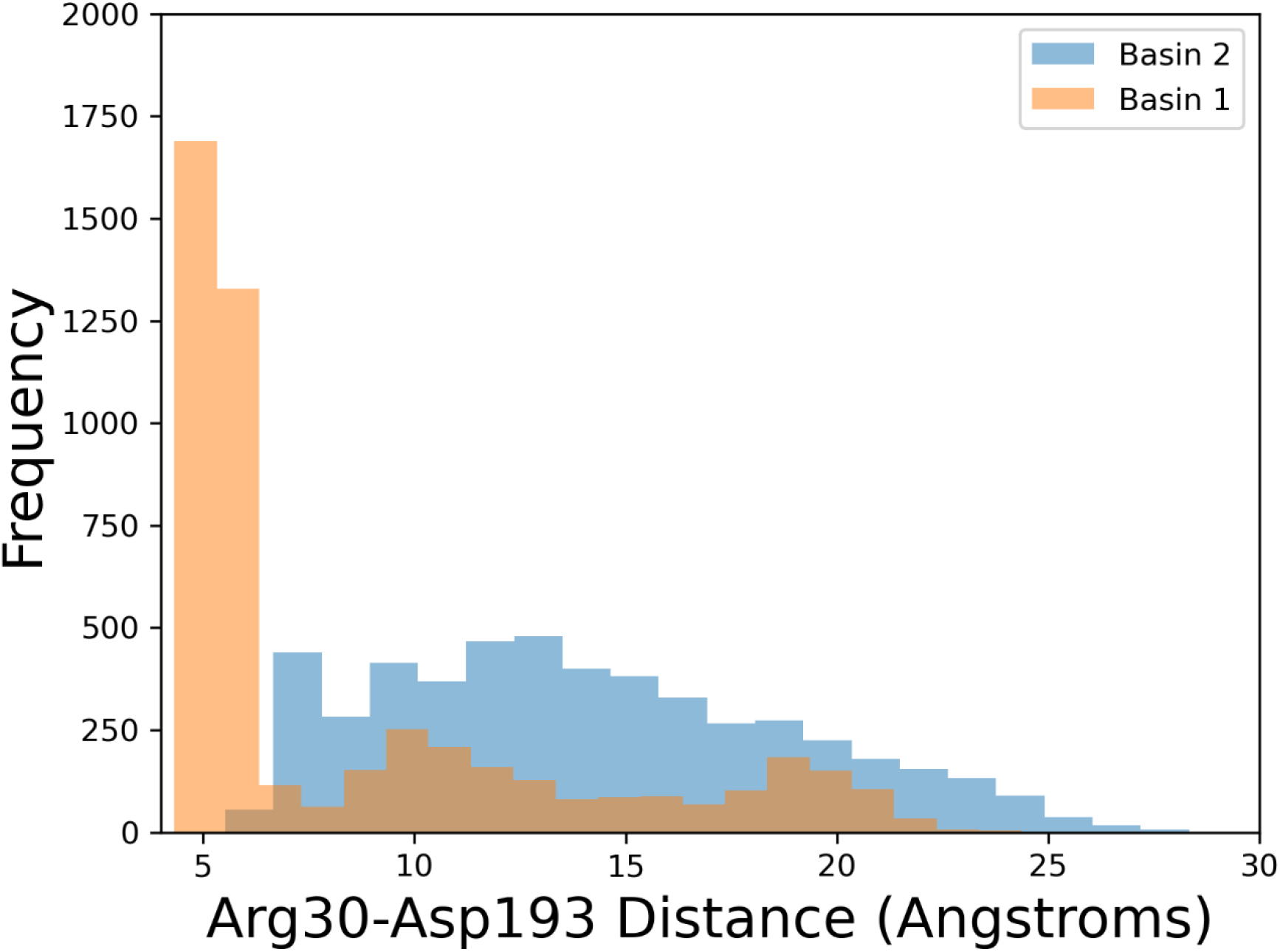
Asp30-Arg193 distance distributions from unbiased MD simulations starting from the centroid structures of basin 1 (orange) and basin 2 (blue) of rMD simulations of PAI-1 in the absence of vitronectin.

**Figure S2.**
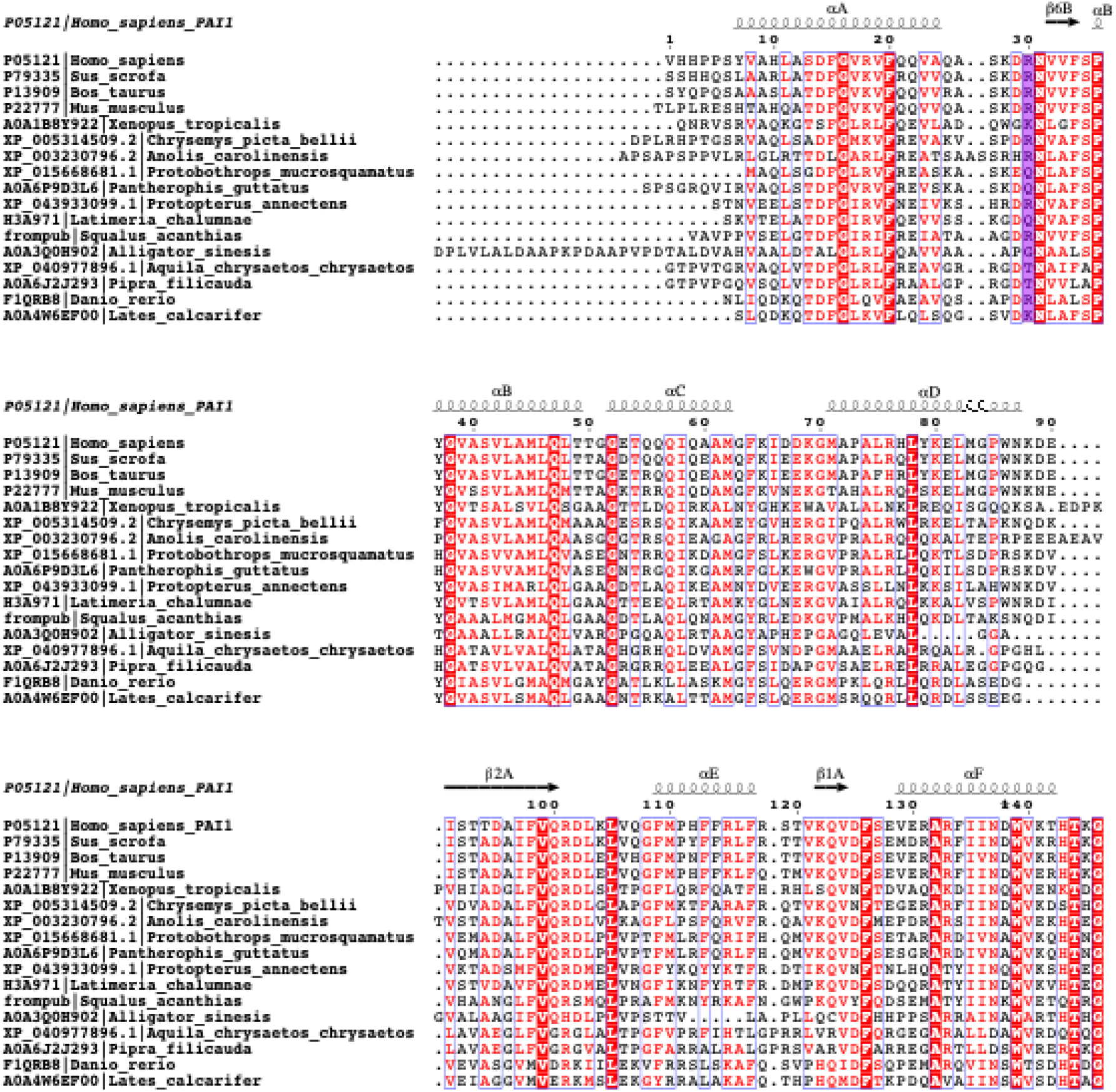
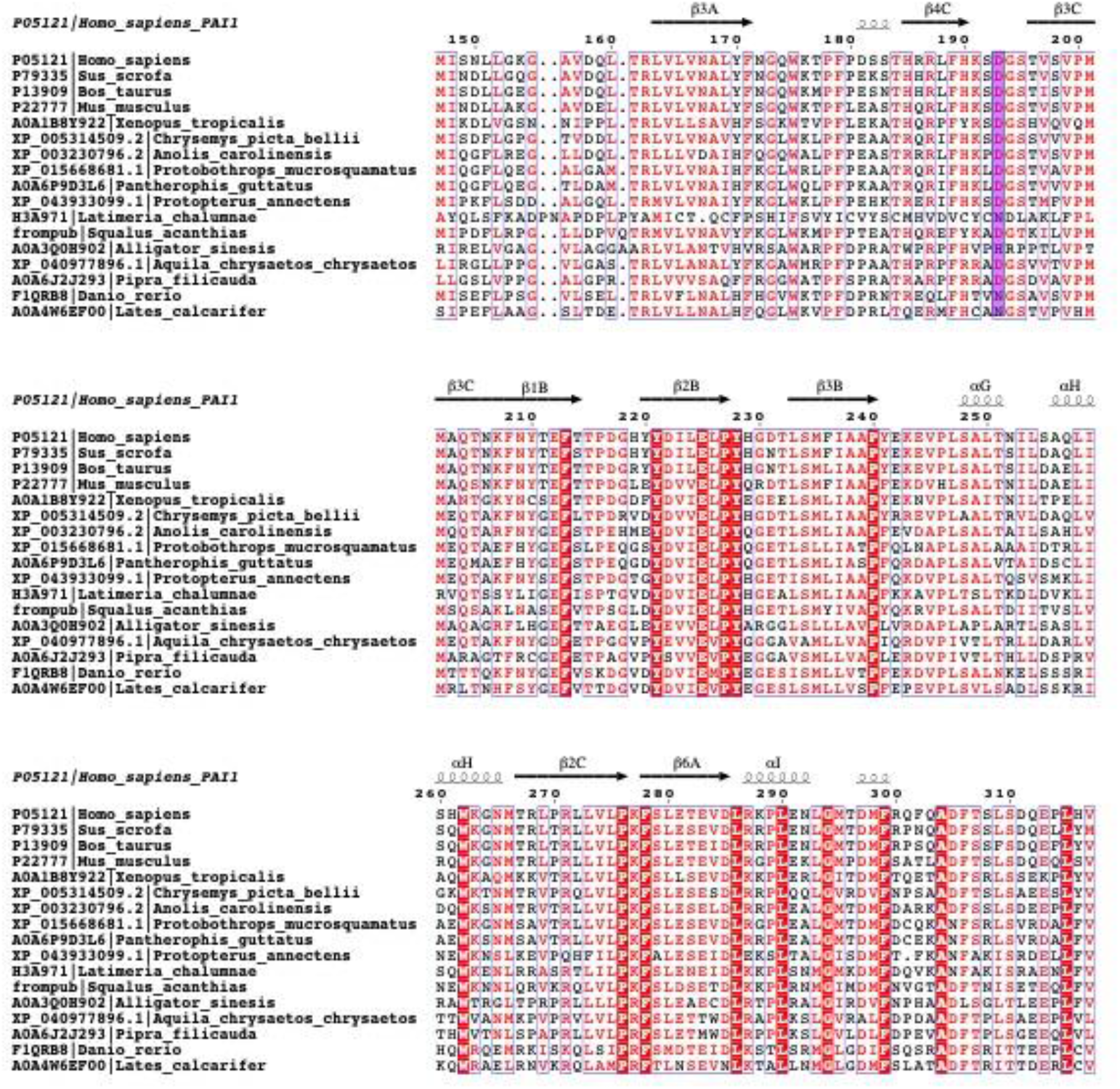
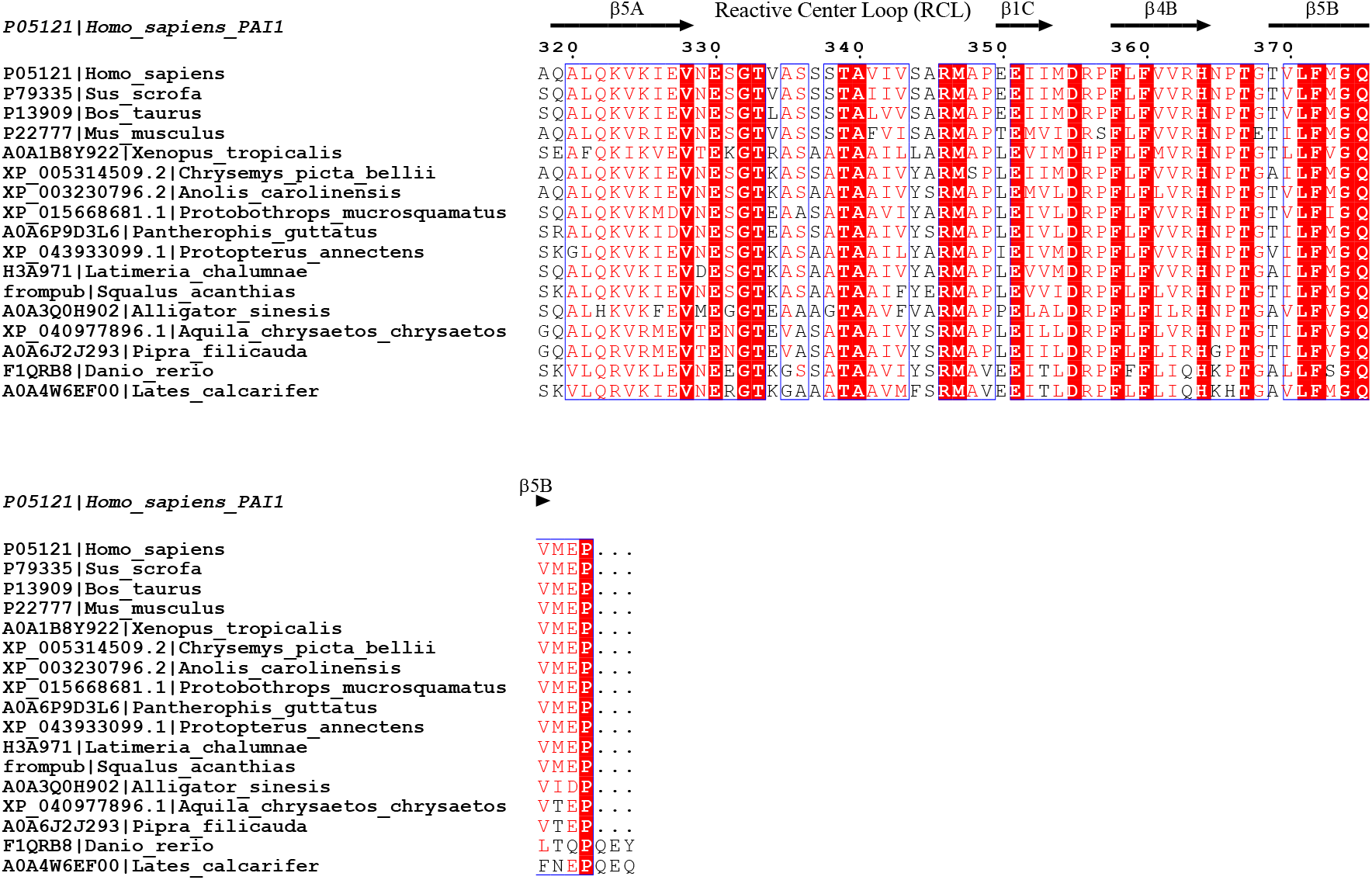
Annotated alignment of 16 PAI-1 sequences from jawed vertebrates. Positions corresponding to Arg30 and Asp193 in human PAI-1 are highlighted in purple.

**Figure S3.**
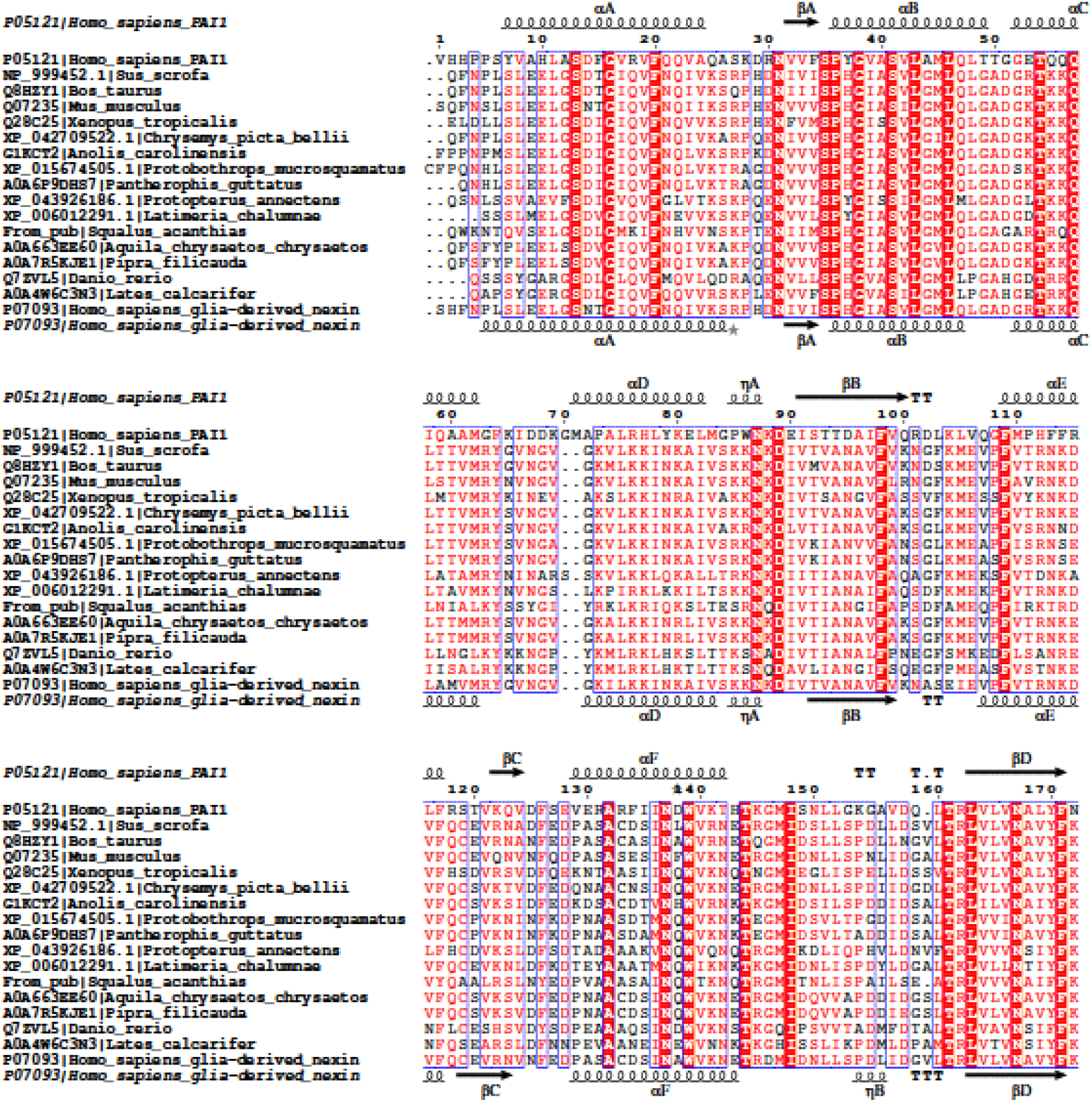
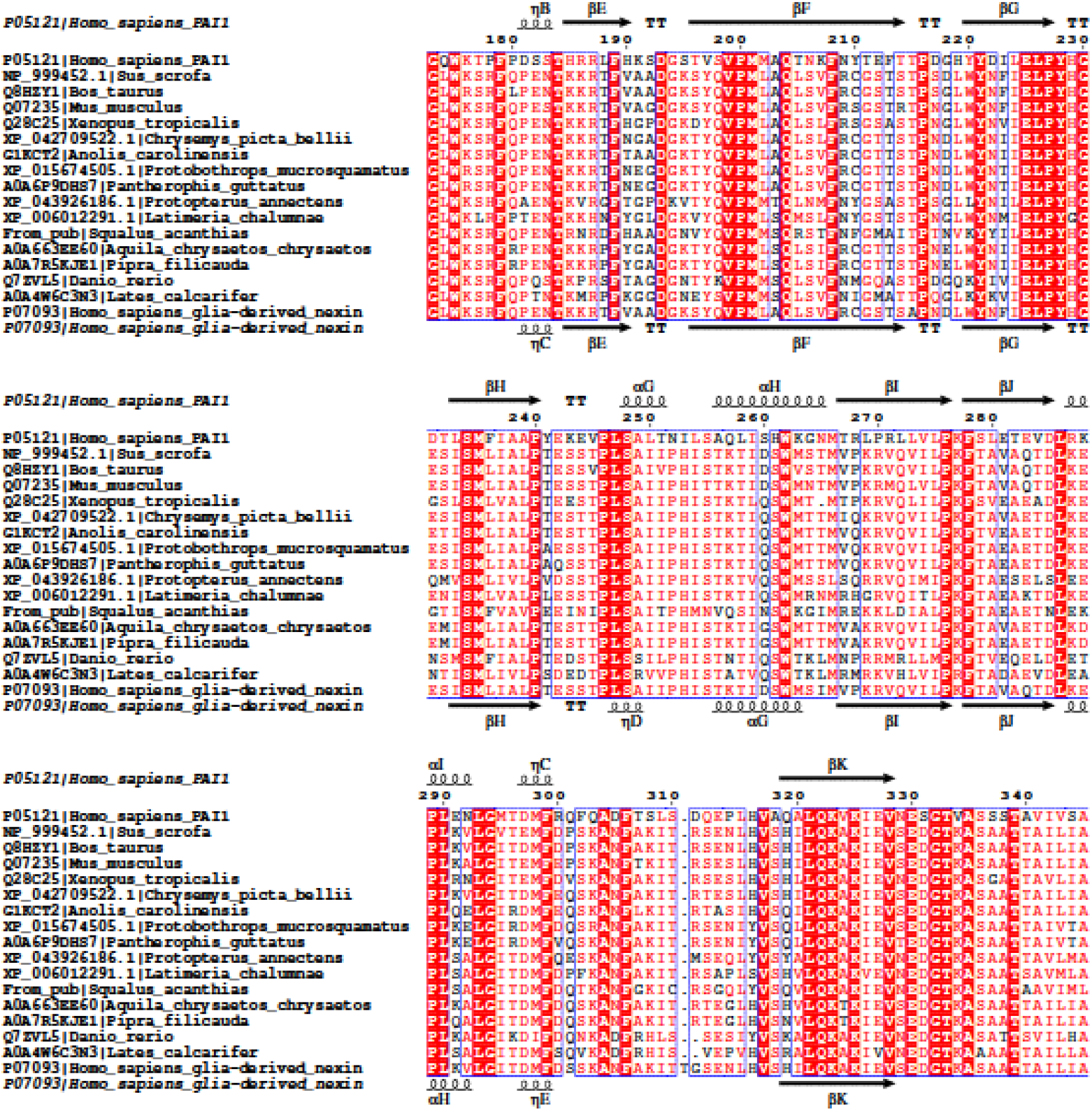
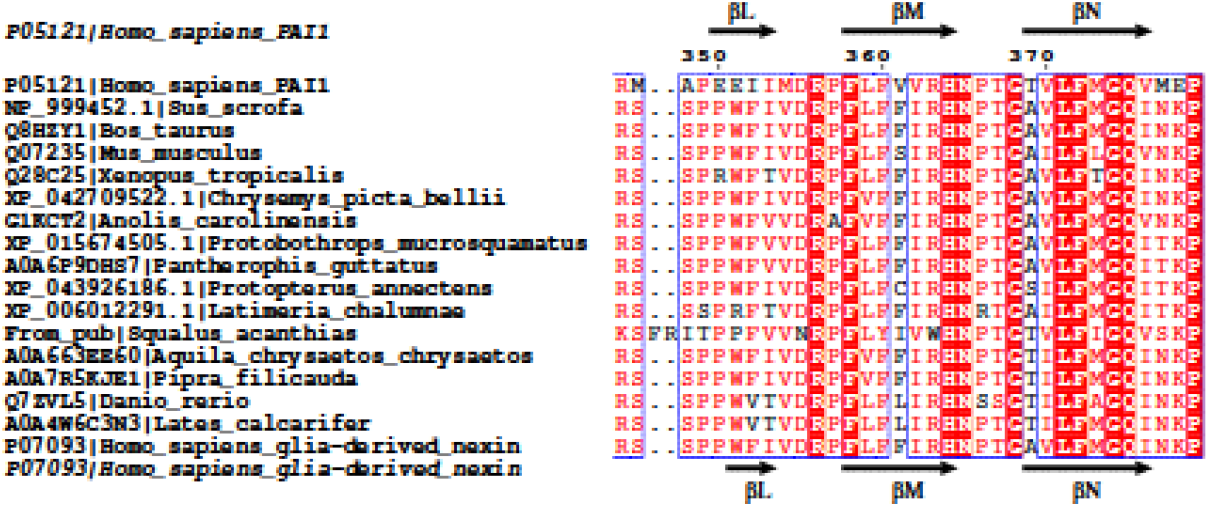
Annotated aligment of human PAI-1 and 16 members of the SerpinE2 family from jawed vertebrates. Positions corresponding to Arg30 and Asp193 in human PAI-1 are highlighted in purple.

